# Identifying sexually dimorphic circulating microRNAs in gonochoristic and hermaphroditic marine fish species

**DOI:** 10.1101/2024.10.11.617774

**Authors:** Benjamin Geffroy, Camille Houdelet, Eva Blondeau-Bidet, Xavier Mialhe, Aline Bajek, Jean-Claude Falguière, Yann Guiguen, Julien Bobe

## Abstract

In many fish species, males and females are hard to distinguish at the juvenile stage and size differences appear during maturation, often favoring females. Molecular tools for sexing live fish would benefit aquaculture, fisheries, and conservation research. Here we aimed to explores to what extent circulating microRNAs (miRNAs) can decipher phenotypic males from phenotypic females in gonochoristic fish species (the European seabass *Dicentrarchus labrax*, the Turbot *Scophthalmus maximus*, the Red drum *Sciaenops ocellatus*, and the Blue Runner *Caranx crysos*) as well as sexual stages in a protandrous hermaphrodite species (the Gilthead seabream *Sparus aurata*). We collected and extracted total RNA from 258 plasma samples, of which 96 samples with satisfactory RNA quality were sequenced using small RNA-seq. Circulating miRNAs detected in the plasma allowed to easily discriminate species, but some miRNAs were significantly correlated to one sex, independently of the species. In immature fishes, 3 miRNAs: miR-21a-3p, miR-18a-3p, and miR-29-1a-5p were overexpressed in females compared to males, while in mature fish, miR-21a-3p exhibited an opposite pattern. In the Gilthead seabream, we detected that both the miR-21a-3p and the miR-125b-2/3-5p were likely involved in the sexual transition from male to female. A complementary analysis on the 3′UTR sequences of all fish species allowed to predict potential mRNA targets of those two miRNAs, some of them being particularly relevant regarding sexual development (*i*.*e*. wnt4, esrrb, esrrga and hsd17b1). The identification of miRNAs like miR-21a-3p and miR-125b-2/3-5p as potential sex markers could offer a new, poorly-invasive method to monitor sex and developmental stages in fishes.

## 1. Introduction

In many fish species, it is almost impossible to distinguish males from females based on external phenotypic characteristics such as color or size at the juvenile stage. Sexual dimorphism, when it appears, is often materialized by differences in size that occurs during maturation. For most marine fishes, the sexual size dimorphism is in favor of females, although this general pattern is not the rule (Horne et al., 2020). Having access to molecular tools allowing to sex any alive fish would thus be a great asset for both aquaculture and fisheries sectors, but also for research with conservation purpose. For instance, in commonly produced marine fish species such as the European seabass *Dicentrarchus labrax*, the Gilthead seabream *Sparus aurata*, the Turbot *Scophthalmus maximus* and the Red drum *Sciaenops ocellatus*, the sex of individuals can only be accurately assessed during maturation, when it is possible to collect gametes. This poses a problem for broodstock management. First, late maturation complicates breeding programs by requiring to keep individuals for many years, even though they may potentially not be used. Second, in the above-mentioned fishes, mature females are much bigger than males, and producing predominantly females would be economically relevant for the aquaculture industry. Having reliable information of sex would also benefit research on wild fish movements patterns such as for the Blue Runner *Caranx crysos* (Brown et al., 2010) that are impossible to sex through external characteristics. This will hence participate to increase our knowledge on fish reproduction and other life history traits by working on alive fish, not on fish purposely sacrificed to know their sex (Bakari et al., 2024; Sley et al., 2012). Finally, non-lethal samplings would also participate to the “3Rs rule” which aim at Replacing, Reducing (the number of animals used), and Refining (anesthetize, provide pain relief) the use of animals for scientific purpose in the context of increasing their welfare.

The development of highly reliable alternatives to hormonal measurements that would be easy-to-implement and require low sample volume would therefore be of great interest. Circulating microRNAs (miRNAs) in the plasma as markers of sex in fish perfectly fulfill these roles. MiRNAs, short and conserved non-coding nucleotide sequences typically spanning 20-22 nucleotides, play pivotal roles in regulating numerous biological processes, mainly by post-transcriptionally repressing target mRNAs (Bartel, 2018). These microRNAs function as gene regulators through base-pairing interactions with mRNAs, with a sequence similarity as low as 6 to 8 nucleotides (seed sequence) being adequate to disrupt Mrna function. Consequently, a single miRNA has the potential to interact with multiple mRNAs, potentially numbering in the hundreds, while individual mRNAs may also be targeted by multiple miRNAs (Bartel, 2009). MiRNAs exhibit high evolutionary conservation, supporting the notion of their fundamental role in metazoan biology (Bartel, 2018). They can also be quantified in various organs but also in extracellular fluids such as serum, blood plasma, urine, and saliva (Mohr and Mott, 2015). These advantageous characteristics of miRNAs make them highly suitable as biomarkers for tracking physiological states. Since their discovery, miRNAs have been extensively employed in human health diagnostics, particularly for tumor detection (Duttagupta et al., 2011; Lu et al., 2005), with growing interest in their application as biomarkers across other species, including fishes (Cardona et al., 2021, 2022). Evidence suggests distinct expression patterns of miRNAs between male and female gonads in diverse fish species (Bhat et al., 2020; Gu et al., 2014; Jing et al., 2014; Tao et al., 2016). However, until now, sex-specific miRNAs have not yet been identified in extracellular fluids such as plasma.

Here, we propose to identify potential circulating miRNAs as sex markers in the plasma of both immature and mature fish belonging to four gonochoristic species: the European seabass, the Turbot, the Blue runner and the Red drum, as well as one protandrous hermaphroditic species, the Gilthead seabream that naturally changes sex from male to female. To do so, we performed a deep small-RNA sequencing on a large number of blood plasma samples of the above-mentioned fishes.

## 2. Material and methods

Experiments were performed in accordance with relevant guidelines and regulations provided by the ethic committee (no 36) of the French Ministry of Higher Education, Research and Innovation and the experiment received the following agreement number: APAFIS #30612-2021031812193539.

### 2.1. Sample collection

All samples were collected between 2020 and 2021. Regarding European Seabass, a total of forty immature (mean weight: 44 ± 11 g and length: 15 ± 1 cm) and twenty mature (1387 ± 303 g and 46 ± 4 cm) fish were randomly collected from the experimental aquaculture station of Ifremer (Palavas-les-Flots, France) and Aquanord aquaculture site (Gravelines, France) respectively. Sixty-five Gilthead Seabream at different reproductive stages (see sex assessment session below and Table 2) were randomly collected from Aquanord aquaculture site (Gravelines, France). Thirty-one wild adult Blue Runners (mean weight 303 ± 70 g and length 24 ± 1.7 cm) were collected at the Station Ifremer Le Robert (Martinique, France) as well as forty-two hatchery produced red drum (mean weight 478 ± 102 g and length 34 ± 2.3 cm). Thirty-two immature turbots (mean weight 110 ± 20 g and length 19 ± 1.2 cm) and twenty-eight immature turbots (mean weight 880 ± 240 g and length 35 ± 2.9 cm) were collected at the France Turbot aquaculture site (Tredarzec, France).

All fish were euthanized with a lethal dose benzocaine (150mg/L). For each individual, a blood sample (1 ml) was collected from the caudal vein thanks to EDTA-coated syringes fitted with 23 G needles. Mature gonochoristic fish were visually sexed, while immature fish were histologically sexed. Regarding the protandrous Gilthead Seabream, a picture of the gonad was taken right after blood sampling, and 5 different stages (from juveniles’ males to adult female) were assessed depending on the ratio of the female/male gonad (Table 1). After centrifugation of blood samples (3000 x g, 5 min), the plasma was stocked at −80°C. Note that the temperature of centrifugation was equal to the temperature at which the fish was living.

**Table 1.** Details of the weight, length, gonadal developmental stages and number of Seabream used for Small RNA sequencing.

|  | n | Weight (g) | Length (cm) | Gonad Ratio Female/Male |
| --- | --- | --- | --- | --- |
| Immature Male | 10 | $221 \pm 45$ | $21 \pm 1$ | |
| Mature Male | 5 | $854 \pm 170$ | $32 \pm 2$ | 1.8 |
| Intersex 1 | 10 | $840 \pm 129$ | $33 \pm 1$ | 18 |
| Intersex 2 | 4 | $870 \pm 179$ | $33 \pm 2$ | 48 |
| Mature Female | 9 | $1745 \pm 244$ | $41 \pm 1$ | |

**Table 2.** Details of the weight, length, maturity stage and number of fish of each species used for Small RNA sequencing.

|  | Blue runner |  |  | Turbot |  |  |  |  |  |
| --- | --- | --- | --- | --- | --- | --- | --- | --- | --- |
|  | Mature |  |  | Immature |  |  | Mature |  |  |
|  | n | Weight (g) | Length (cm) | n | Weight (g) | Length (cm) | n | Weight (g) | Length (cm) |
| Females | 3 | 281 ± 33 | 24 ± 0.9 | 8 | 110 ± 25 | 19 ± 2 | 5 | 935 ± 250 | 35.5 ± 3 |
| Males | 3 | 320 ± 77 | 24 ± 2 | 10 | 119 ± 15 | 20 ± 1 | 3 | 766 ± 232 | 34 ± 4 |
|  | Red Drum |  |  | European Seabass |  |  |  |  |  |
|  | Immature |  |  | Immature |  |  | Mature |  |  |
|  | n | Weight (g) | Length (cm) | n | Weight (g) | Length (cm) | n | Weight (g) | Length (cm) |
| Females | 5 | 508 ± 109 | 35 ± 2 | 3 | 34 ± 2 | 14 ± 0.1 | 3 | 957 ± 161 | 45 ± 7 |
| Males | 8 | 516 ± 75 | 35 ± 2 | 4 | 45 ± 13 | 15 ± 1 | 3 | 1331 ± 138 | 47 ± 1 |

### Identification of markers

#### Histology

Gonads of immature fishes (European seabass, Turbot and Red Drum) were fixed in Bouin’s fluid for 6 to 8 h and rinsed in clear water for one hour. Then, they were rinsed in EtOH 70% for several days and placed in a dehydration automate (STP 120, MM, France). Each gonad was embedded in paraffin and cut at 5µm sections. Slides were stained using the Masson’s trichrome methods (MYREVA SS30, MM, France).

#### RNA extractions

Thawed plasma of each fish was homogenized in QIAzol lysis reagent (Beverly, MA, USA) following manufacturer’s instructions. The total RNA was resuspended in 15 µL of RNAse free water. MiRNAs were quantified using the smallRNA Analysis kit (DNF-470-0275) on a Fragment Analyzer (Agilent). Based on the fragmentation profile (with or without pic between 17 and 25 nucleotides), the estimated quantity (ng/μl) and percentage (%) of miRNAs relative to other small RNAs, a subsample of individuals was selected for libraries’ construction.

#### Small RNA Sequencing

Libraries were constructed using the NEXTFLEX Small RNA-seq kit v3 (Perkin Elmer, #NOVA-5132-05). Briefly, a 3’ Adenylated adapter was ligated to the 3’ end of 0,5 ng of microRNA and purified to remove 3’ adapter excess. A 5’ adapter was ligated to the 5’ end of the 3’ ligated microRNA. The resulting construction was purified to remove 5’ adapter excess. 5’ and 3’ ligated microRNAs underwent a reverse transcription using a M-MuLV reverse Transcriptase and a RT primer complementary to the 3’ adapter. Resulting cDNAs were used as a matrix in a 25 cycles PCR using a pair of uniquely barcoded primers. The resulting barcoded library was size selected on a Pippin HT (SAGE Science) using 3 % agarose cassette (#HTG3004) and aiming for a size range between 147 bp and 193 bp. Once size selected, libraries were verified on a Fragment Analyzer using the High Sensitivity NGS kit (#DNF-474-0500) and quantified using the KAPA Library quantification kit (Roche, ref. KK4824). Following this second quality check, we selected 38 samples of Seabream (Table 1), 6 samples of Blue Runners, 13 samples of Red Drum, 26 samples of Turbots and 13 samples of European sea bass (Table 2) for being sequenced.

#### MiRNA alignment and quantification

Image analyses and base calling were performed using the NovaSeq Control Software and Real-Time Analysis component (Illumina). Demultiplexing was performed using Illumina’s conversion software (bcl2fastq 2.20). The quality of the raw data was assessed using FastQC from the Babraham Institute and the Illumina software SAV (Sequencing Analysis Viewer). The raw reads were trimmed using Cutadapt (version 3.5) (Martin, 2011) to remove the sequencing adapter (TGGAATTCTCGGGTGCCAAGG) at the 3′-end. Additionally, 4 bases were also trimmed from the 5′-end and 3′-end of the reads as indicated in the manual of NEXTflex Small RNA-Seq Kit v3 from Bio Scientific. Before counting step, samples with a rRNA degradation profile were filtered out. This resulted in 13 and 16 immature and mature fish, respectively, for the experiment on sex and 21 fish for the experiment on stress. MiRNA analysis was performed with Prost ! v0.7.60 pipeline (Desvignes et al., 2019). In this pipeline, alignment was performed with BBMap (Bushnell, 2014) version 38.90 to genome of interest : the European sea bass *Dicentrarchus labrax* GCA_000689215.1_seabass_V1.0, the one of the Red drum *Sciaenops ocellatus* (GCA_014183145.1_ASM1418314v1), the Gilthead seabream *Sparus aurata* (GCF_900880675.1_fSpaAur1.1_genomic.fna), the Jack C*aranx ignobilis*, phylogenetically closest species of the Blue Runner *Caranx crysos* (GCA_021432705.1) and the Turbot *Scophthalmus maximus* (GCF_013347765.1). Next, sequences with overlapping genomic positions were grouped together. Only sequences with a minimum of 30 reads (when adding up all samples for a given species) were retained, to perform the differential expression analysis. The miRNA annotation and counting were retrieved by Prost! (parameters in Text S1) from a custom annotation provided in FishmiRNA (https://www.fishmirna.org/) using *Gasterosteus aculeatus* miRNA sequences (the phylogenetically closest species).

#### Statistical analysis

A Non-metric Multi-dimensional Scaling (NMDS) approach was used to compare species, sex and gonadal maturity stages based on all miRNAs sequenced. A r^2^ analysis indicated a significant effect according to the condition tested. The package vegan v2.6-2 (Oksanen et al., 2013) was used and the construction of the dissimilarity matrix was based on the Bray-Curtis methods.

Differentially expressed miRNA were identified using one Bioconductor (Gentleman et al., 2004) package: DESeq2 1.32.0 (Love et al., 2014). Data were normalized using the default method for DESeq2 package. MiRNA with adjusted p-value below 5% (according to the FDR method from Benjamini-Hochberg) were considered significantly differentially expressed between sex (males or females) or gonadal stages for the Gilthead seabream. To better depict the miRNAs potentially discriminating males from females we conducted a Principal Component Analysis (PCA) on miRNAs with unadjusted p-value below 5% following the DESeq2 comparing all species together for only immature or mature stages. We also performed a NMDS to discriminate gonadal stages over sex-change of the Gilthead seabream. All statistical analyses were conducted with the R v 4.1.0 (Core Team, 2020).

### Prediction of target

We retrieved the 3’UTR sequences from the European seabass, the Red drum, the Gilthead seabream and the Turbot (note that annotated 3’UTR sequences were not available for the Blue Runner). To identify miRNA targets, we used the freely available Perl script of TargetScanHuman 8.0. (https://www.targetscan.org/vert_80/) (Agarwal et al., 2015) that we adapted for our fish species. This allowed to obtain a list of potential target genes by searching for the presence of conserved 8mer, 7mer, and 6mer sites that match the seed region of each miRNA of interest. Then all sequences of potential target genes for the 4 fish species were aligned using the MUSCLE Alignment in Geneious Prime 2019.2.3 and the position of the seed was confirmed in order to corroborate that the position of the match is evolutionary conserved between these species, enforcing the credibility of the target.

## Results

On the 258 plasma samples collected and for which the RNA was extracted, a total of 96 samples for which the quality of the RNA was considered satisfying were sequenced. The Non-metric multidimensional scaling (NMDS) of all miRNAs detected in the plasma of the four gonochoristic fish species allowed to well discriminate the species (r^2^ = 0.82; p-value < 0.001***; Figure 1); and gonadal developmental stages (mature vs immature, r^2^ = 0.07; p-value < 0.05*) but not sex (females vs males, r^2^ = 0.003; p-value = 0.87). Therefore, subsequent analyses were performed within each gonadal stage.

**Figure 1.**
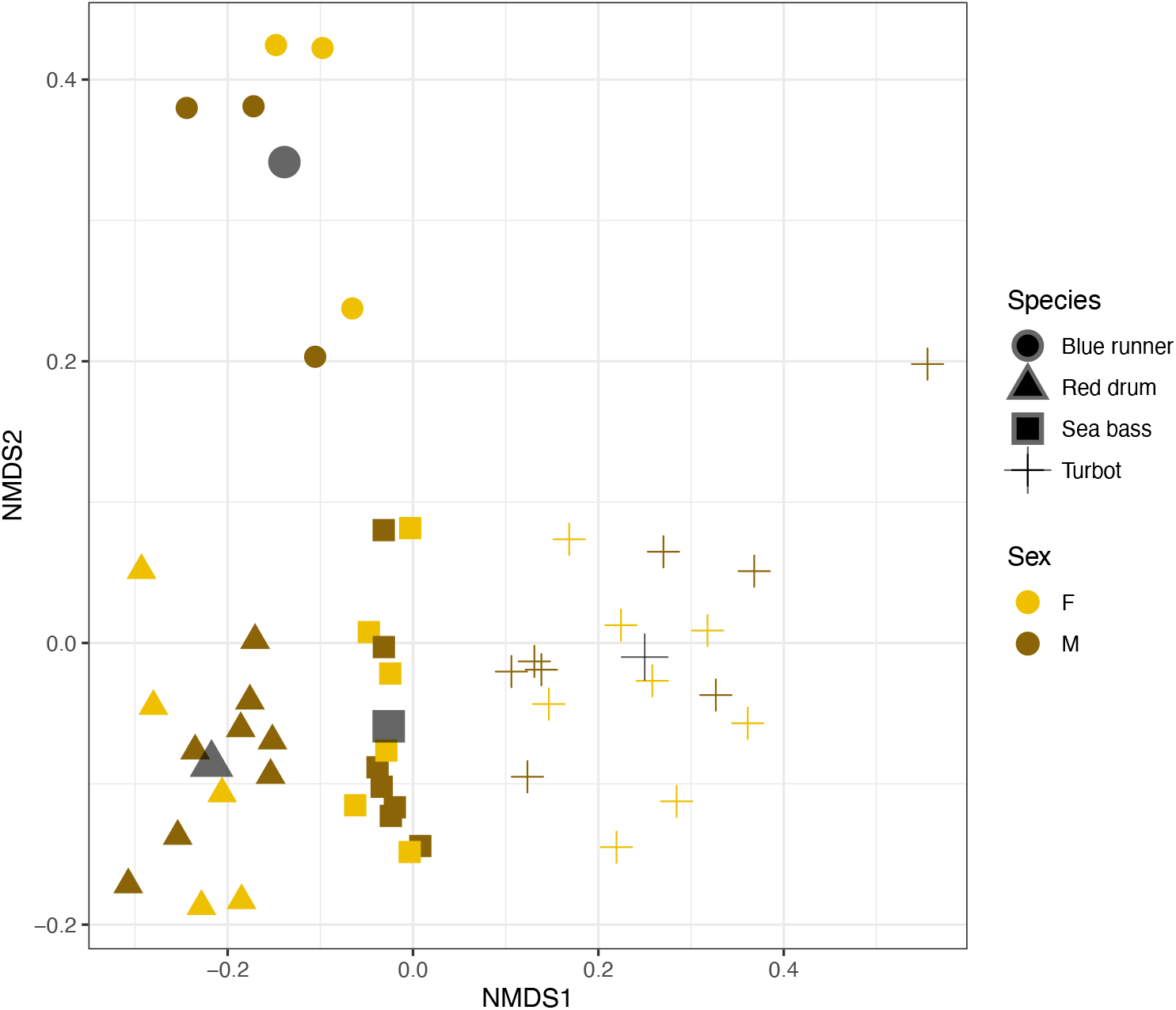
Non-metric multidimensional scaling (NMDS) of all miRNAs detected in the plasma of four gonochoristic fishes. On NMDS, each point represents an individual form each of the 4 species, differing by their shape. Yellow points represent females, while brown points represent males.

### Immature gonochoristic fish

From the DESEQ2 analysis, we detected 19 miRNAs for which uncorrected p-value were below 0.05, that allowed to well discriminate males from females in the first axis of the PCA (46 % of variance explained), independently of the species (Figure 2A). Among those 19 miRNAs 3 were highly significant: miR-21a-3p, miR-18a-3p and miR-29-1a-5p (Figure 2A, in red, corrected p-value < 0.05), although this pattern was mainly driven by Turbots.

**Figure 2.**
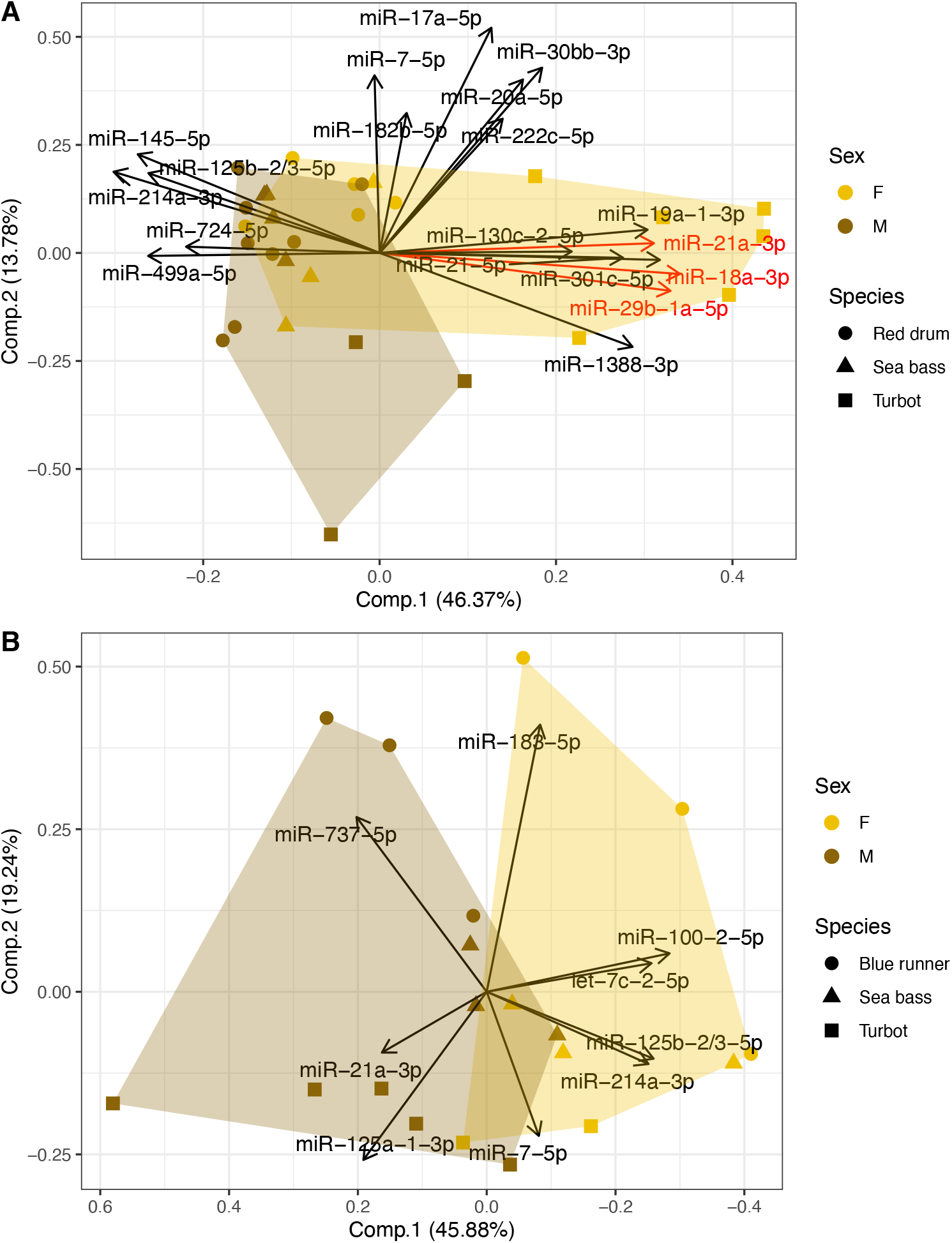
PCA of the most differentially expressed miRNAs in the plasma of A) Immature gonochoristic fish and B) Mature gonochoristic fish. Each symbol represent an individual of a species, and each arrow, a variable : miRNA. MiRNAs represented in red are significant (corrected p-value < 0.05) in the DeSeq2 analysis. Females and males are represented by yellow and brown dots respectively.

### Mature gonochoristic fish

The DESEQ2 analysis yielded 9 miRNAs for which uncorrected p-values were below 0.05, that allowed to well discriminate mature males from mature females in the first axis of the PCA (46 % of variance explained), independently of the species (Figure 2B). None of them were significant when applying the FDR correction method from Benjamini-Hochberg.

Overall, 3 miRNAs were common to both analyses. The miR-7-5p presented a similar pattern between mature and immatures samples, with females tending to have more miR-7-5p than males. On the contrary, both and the miR-125b-2/3-5p and miR-21a-3p presented an opposite pattern between males and females according to the maturity stage. When considering each species, we detected significantly more miR-23a-1-5p in males compared to females as well as significantly more miR-206-1-3p and miR-1-3p in females compared to males in Jacks (adjusted p-values < 0.05 for all comparisons). In immature Red drum we could not identify miRNAs differentially expressed between sex. In the European seabass only, we identified 11 differentially expressed miRNAs between males and females (Houdelet et al. 2023), and two of them were in the common analysis of immature fish, the miR-499a-5p and the miR-1388-3p.

### In a protandrous hermaphroditic fish: The Gilthead seabream

The DESEQ2 analyses allowed to detect a total of 76 unique miRNAs differentially expressed between all stages (Supplementary Data 1). Within these 76 miRNAs, 54 miRNAs were at least common to two comparisons (e.g. between immature male and mature male as well as between immature male and intersex 1). The more prominent comparison was between immature males and mature females, with 34 differentially expressed miRNAs (Figure 3A). The NMDS of all miRNAs (stress = 0.16) detected in the plasma allowed to significantly distinguish the five different stages (r^2^ = 0.28; p-value = 0.01; Figure 3A, 3B). When focusing on all those discriminating significantly all stages in a pair-wise comparison (1 stage vs 1 Stage, using DESEQ2), the effect of gonadal stage on this new NMDS (stress = 0.17) was more pronounced (r^2^ = 0.46; p-value < 0.001, Figure 3C). Since the Gilthead seabream change sex at the same time its grow, it was difficult to decipher sex and age-related miRNAs. Interestingly, we found out that two of our markers detected in immature and mature fish, miR-21a-3p and miR-125b-2/3-5p (Figure 2) were also significantly and differentially expressed between both immature and mature males and mature females in the Gilthead seabream (Figure 4,5). The miR-125b-2/3-5p presence was dependent on the developmental stage, with more reads in mature females compared to mature males and more miR-125b-2/3-5p reads in immature males compared to immature females (Figure 5).

**Figure 3.**
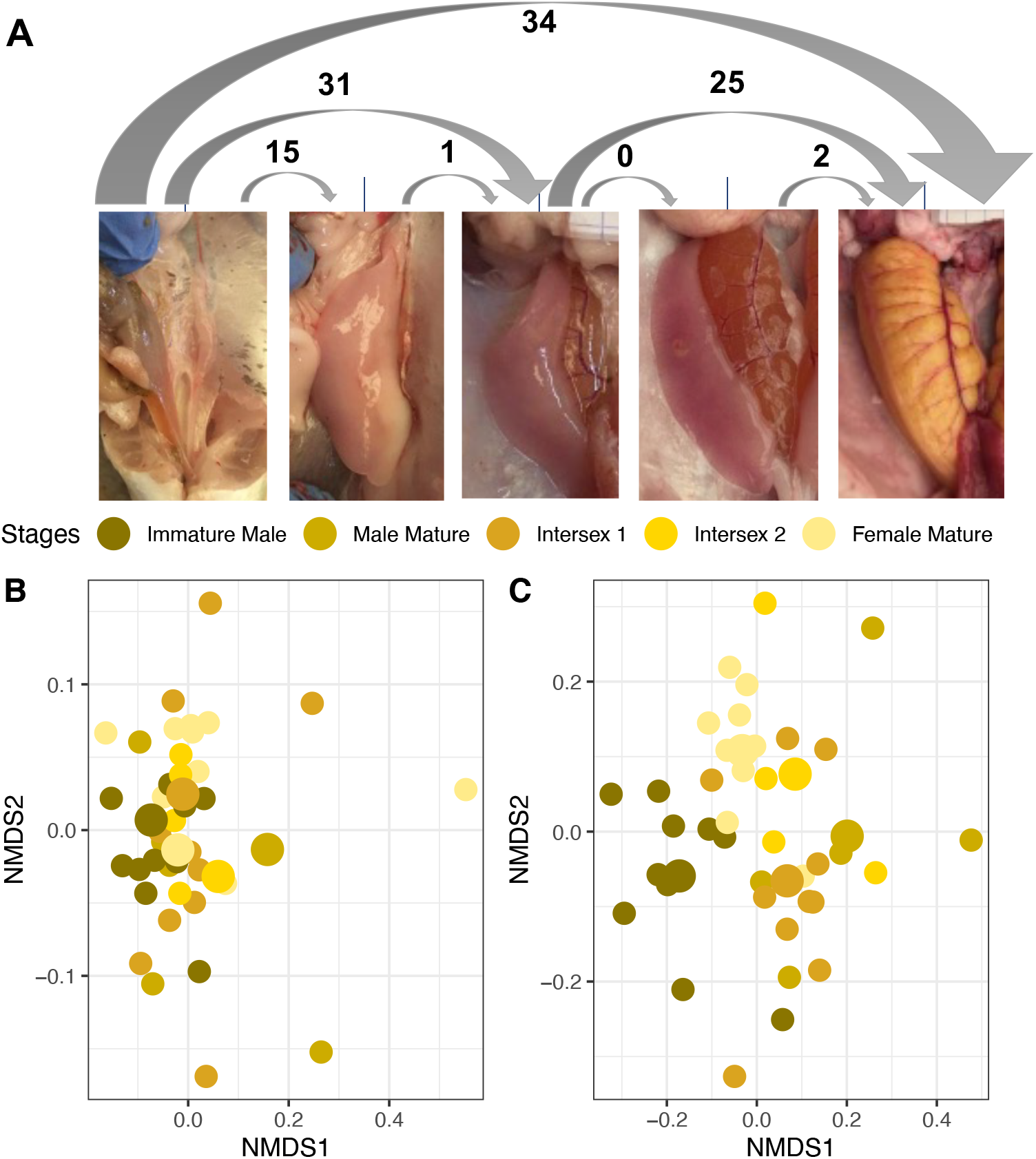
(A) The 5 different gonadal stages, for which plasmatic miRNAs were sequenced. The number bellow each arrow represent the number of miRNAs differentially expressed between stages. (B) Non-metric multidimensional scaling (NMDS) of all miRNAs detected in the plasma of the Gilthead seabream according to their gonadal developmental stages. (C) NMDS of all miRNAs differentially expressed between the different gonadal developmental stages. The different colors distinguish the different developmental stages.

**Figure 4.**
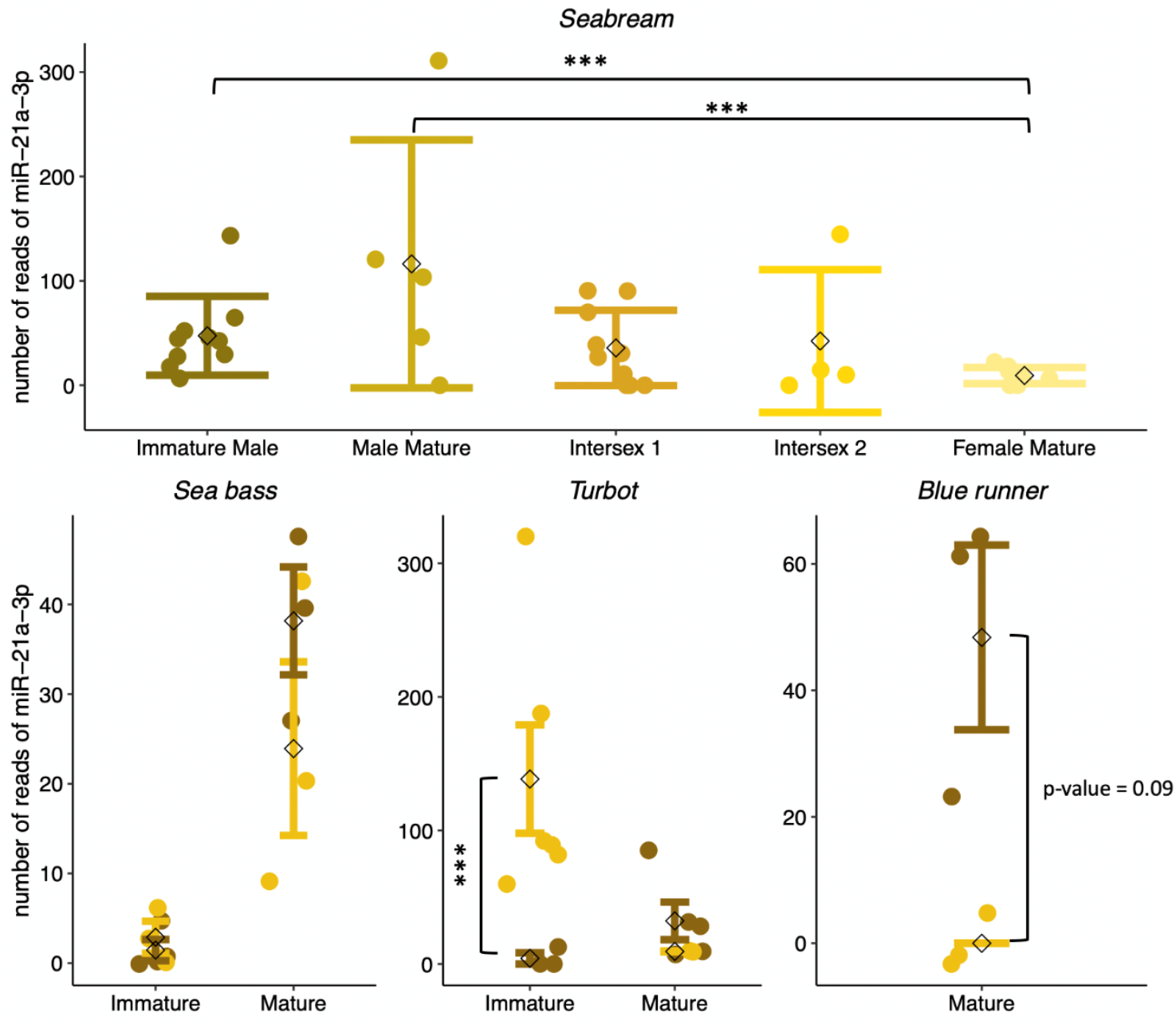
Copy number of miR-21a-3p in Gilthead Seabream, European Sea bass, Turbots and Blue runner. Females and males are represented by yellow and brown dots respectively. Each bar represents the mean (a black diamond) ± se.

**Figure 5.**
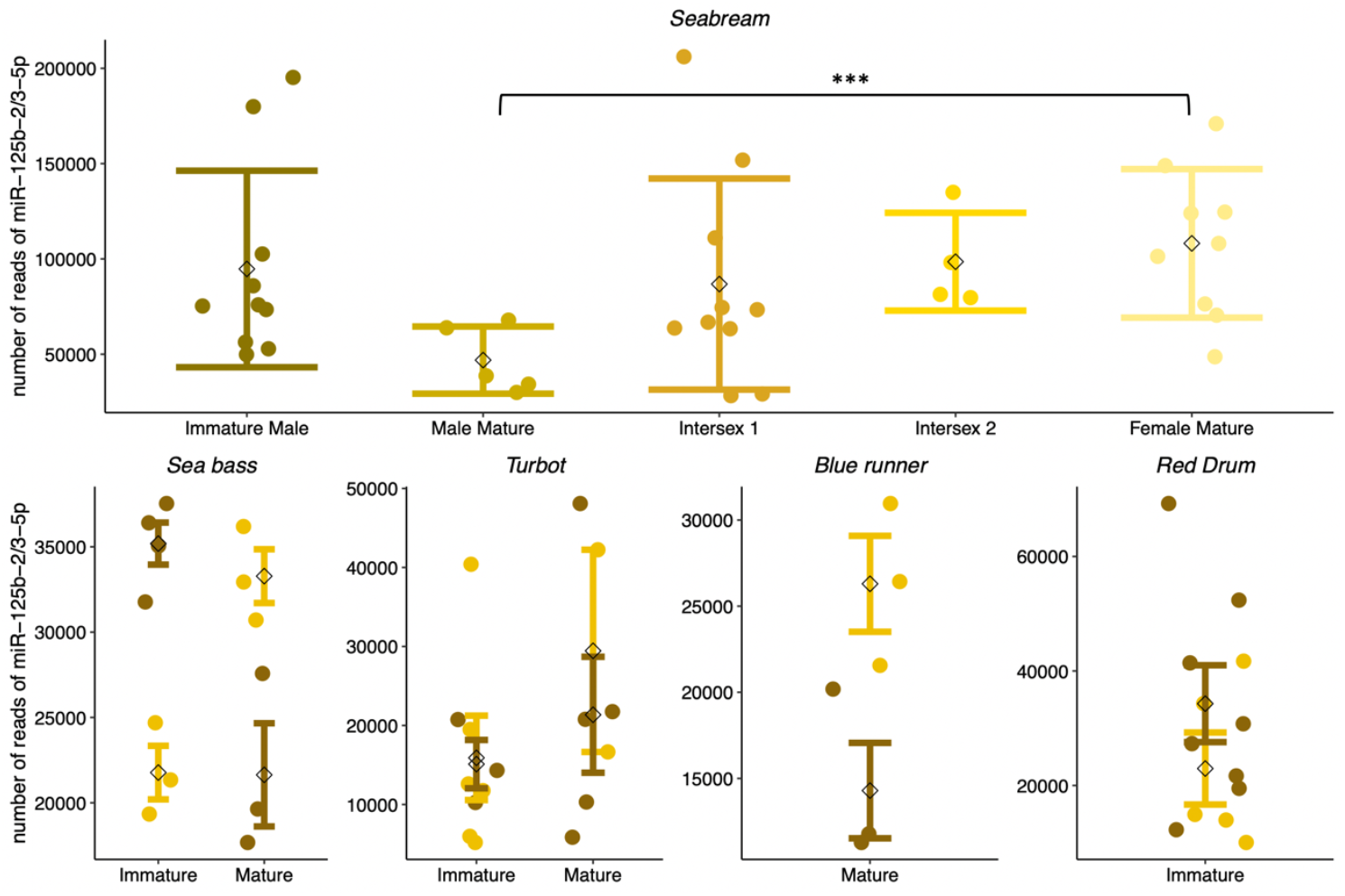
Copy number of miR-125b-2/3-5p in Gilthead Seabream, European Sea bass, Turbots, Blue runner and Red Drum. Females and males are represented by yellow and brown dots respectively. Each bar represents the mean (a black diamond) ± se.

### Target prediction of miR-21a-3p and miR-125b-2/3-5p

The seed sequence used for miR-21a-3p was GACAACA and CCCTGAG for miR-125b-2/3-5p. Those seeds were common for all fish species. We detected 23 possible gene targets of miR-21a-3p that were common to 4 species: the Gilthead Seabream, the European Seabass, the Turbot, and the Red Drum; and 184 targets that were common to at least 3 of these 4 species (Supplementary data 2). Among the 184 genes, 6 were of interested regarding hormonal regulation: wingless-type MMTV integration site family, member 4 (wnt4), estrogen-related receptor beta (esrrb), estrogen-related receptor gamma a (esrrga), insulin-like growth factor 2 mRNA binding protein 2a (igf2bp2a), insulin-like growth factor 1 (igf1) and insulin receptor a (insra). To confirm that the seed region of miR-21a-3p could bind to the same position of the 3’UTR sequence for all fish, they were all aligned using Geneious

Prime and a consensus sequence was built. For wnt4, all 3’URT aligned completely with the 7 bases of the seed region of miR-21a-3p (Supplementary Figure 1). For insra, the 3’UTR sequence of the Turbot was not available, but the seed region of miR-21a-3p aligned at a common 3’UTR sequence region for the Red drum, the European seabass and the Gilthead seabream. For esrrga, the seed region aligned with the 3’UTR of the Red drum, the European seabass and the Turbot. There was only one base difference for the Gilthead seabream (A instead of G; Supplementary Figure 1), which could also be an error of sequencing. For esrrb, the seed region aligned with the 3’UTR of the Gilthead seabream, the European seabass and the Turbot. There was only one base difference for the Red drum (C instead of G; Supplementary Figure 1). Regarding igf1, only 6 bases the seed region of miR-21a-3p aligned with the four 3’UTRs sequences. Regarding igf2bp2a, the seed region aligned with the 3’UTR of the Gilthead seabream, the European seabass and the Red drum at the same location, but not for the Turbot.

We detected 23 possible targets genes of miR-125b-2/3-5p that were common to 4 species: the Gilthead Seabream, the European Seabass, the Turbot, and the Red Drum; and 90 potential targets that were common to at least 3 of these 4 species (Supplementary data 3). Among the 90 genes, 2 were of interested regarding hormonal regulation: Igf2bp2a, which was also a potential target of miR-125b-2/3-5p, although the seed sequence aligned only with 6 nucleotides (6mer) in the four fish species. Another potentially interesting target of miR-125b-2/3-5p was hsd17b1, but it also aligned with only 6 nucleotides at the same region for 3 species: the Gilthead seabream, the European seabass and the Red drum (the 3’UTR sequence of hsd17b1 was not available for the Turbot).

## Discussion

Our study revealed that circulating miRNAs detected in the plasma are highly species-specific, but some miRNAs were significantly correlated to one sex, independent of the species. This is the case for miR-21a-3p, miR-18a-3p, and miR-29-1a-5p, which are over-expressed in immature females compared to immature males. For mature fishes, 9 miRNAs allowed discrimination between males and females, but when tested individually, none of them was significantly associated with one sex (using DESeq2 analysis). Surprisingly, for the gonochoristic fish species for which both immature and mature fish were available (Turbots, European seabass, and Red drum), libraries were successfully constructed for 57% of the RNAs available in immature fish and only for 30% of the RNAs available in mature fish. For instance, 39 RNA samples were available for mature Red drum, but none of them was of sufficient quality for constructing small RNA-seq libraries. This highlight sharp differences in plasma composition between immature and mature fish and might be a reason why we could not detect miRNAs differentially expressed according to sex in mature individuals. Nonetheless, when considering all miRNAs with an uncorrected p-value < 0.05, two miRNAs, namely miR-21a-3p and miR-125b-2/3-5p, were of particular interest with opposite expression patterns between males and females according to the maturity stage. Interestingly, these two miRNAs were also detected as differentially expressed between individuals with a mature testis and those with a mature ovary in the protandrous Gilthead Seabream.

Specifically, miR-21a-3p was over-expressed in Gilthead Seabream presenting a mature testis compared to individuals exhibiting a mature ovary, and the same tendency was observed for mature individuals in gonochoristic species: the European Seabass, the Turbot, and the Blue Runner. The opposite pattern was detected in immature European Seabass and Turbot, where immature females (all species together) presented significantly more reads of miR-21a-3p than immature males. Interestingly, this miRNA was found to be abundantly expressed in ovaries of zebrafish and common carp (*Cyprinus carpio*), depending on the developmental stage (F. Wang et al., 2017; Wong et al., 2018; Zayed et al., 2019). Now, when comparing miRNAs between sex, expression levels of miR-21-3p was more than 4-fold in XX ovary as compared to XY testis of immature (1-year old) yellow catfish (*Pelteobagrus fulvidraco*)(Jing et al., 2014). It is worth noting that none of these studies were performed in plasma, where one could expect lower miRNA quantities relative to organs. For instance, in a pre-experiment in which we sequenced gonads and plasma of the same European seabass individuals (n = 10), we detected about 15 times more miR-21a-3p in the gonad than in the plasma (mean number of reads: 473 vs. 32).

Yet, most of our knowledge regarding the role of miR-21a-3p in controlling gonadal development comes from mammals, such as mice, sheep, porcine, and bovine (Arefnezhad et al., 2024; Reza et al., 2019). MiR-21 plays a significant role in oocyte maturation, as well as in the development of blastocysts and embryos. It is notably overexpressed during the transition from germinal vesicle to oocytes, which are arrested at the metaphase II (Arefnezhad et al., 2024). In addition, MiR-21-3p is highly expressed in atretic follicles compared to healthy follicles (Donadeu et al., 2017). This would corroborate our findings, where immature females (before the start of vitellogenesis) present more copies of this miRNA than immature males in gonochoristic species, and mature males present more miR-21a-3p than mature females. In addition, those results also well align with what we detected in the protandrous Gilthead seabream that possess an ovotestis, and where ovarian development is blocked in mature males, with potentially high numbers of atretic follicles. The *in silico* analysis revealed that miR-21-3p is a potential regulator of numerous signaling pathways, including meiosis, wingless-type MMTV integration site family, member 4 (wnt4), estrogen-related receptors (beta, esrrb and gamma, esrrga), as well as insulin (igf2bp2a, igf1, insra), with conserved target sites among the different fish species investigated. All of these genes play key roles in sex determination and sexual development.

The miR-125b-2/3-5p also presented an interesting expression pattern that sharply differs according to the developmental stage, but in a direction opposite to that of miR-21a-3p. In all fish investigated, females at a mature stage presented more miR-125b-2/3-5p than males, while immature males tended to have more miR-125b-2/3-5p in the plasma than their female counterparts. When comparing stages within sex, these results are not in line with those of Papadaki et al. (2020) who detected more miR-125c (that share the same seed region that miR-125b) in immature females compared to mature females. The authors proposed the follicle-stimulating hormone receptor (FSHR) as a putative target of miR-125c applying RNAhybrid (version 2.12 (Krüger and Rehmsmeier 2006)) (Papadaki et al., 2020), but we were not able to detect such a target for miR-125b-2/3-5p using TargetScan. One interesting putative target concerning sexual development was hsd17b1, an enzyme that catalyzes the final step of estrogen biosynthesis, by preferentially reducing estrone to yield the potent estrogen 17β-estradiol. In female mice, miR-125b expression was found to be activated by estrogen via Erα, both *in vitro* and *in vivo* (Z.-C. Zhang et al., 2015), while the Luteinizing hormone (LH) inhibited miR-125b-5p expression (X. Zhang et al., 2020). Furthermore, inhibiting miR-125b-5p led to an increase in the expression of genes related to androgen synthesis and stimulated testosterone secretion, while simultaneously downregulating genes related to estrogen synthesis and decreasing estradiol release (X. Zhang et al., 2020). These authors identified Pak3 as a direct target of miR-125b-5p, but such a target was not found in our search of putative targets (X. Zhang et al., 2020). Overall, these results highlight an important crosstalk between miR-125b-5p and oestrogen synthesis-related enzymes, while much less is known regarding androgens. Yet, in the half smooth tongue sole (*Cynoglossus semilaevis*), high level of miR-125b-5p were detected in the sperm of fish compared to their exosomes in the seminal plasma (Zhao et al., 2023) and it was found to be one of the most abundantly expressed miRNA in both gonads of the yellow catfish (*Pelteobagrus fulvidraco*)(P. Wang et al., 2018).

Our study identified several miRNAs as potential sex markers for marine fish. However, their expression in plasma is closely linked to the animal’s sexual developmental stage. This is evidenced by the gradual increase or decrease of miR-125b-2/3-5p and miR-21a-3p, respectively, which characterize the transition from male to female in the Gilthead Seabream. Including both gonochoristic and hermaphroditic species in the study helped to eliminate age as a confounding factor in detecting sex-specific miRNAs in the protandrous Gilthead Seabream. Moreover, detecting these miRNAs in plasma is advantageous for monitoring sexual stage transitions and determining sex in a non-invasive manner. We hope our study will pave the way in developing new miRNA-based sensors that can provide insights into sex and sexual developmental stages in fish.

## Data, script, code, and supplementary information availability

All data generated or analyzed during this study are included in this article and in supplementary information. The R script and processed data from the small RNA-seq are available in Zenodo: https://doi.org/10.5281/zenodo.14013730. The sequencing raw data are available through the online ENA (European Nucleotide Archive) platform, under the access number: PRJEB80705.

## Acknowledgments

The authors would like to acknowledge the team from Station Ifremer Palavas-les-Flots (France) and the team from Gloria Maris (France) for the help in different sampling.

