## Supplementary figures and images for "Identifying sexually dimorphic circulating microRNAs in gonochoristic and hermaphroditic marine fish species"

### Supplementary Figure 1

Alignment mir-21a-3p

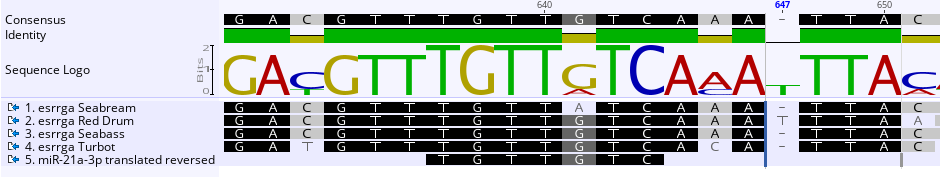

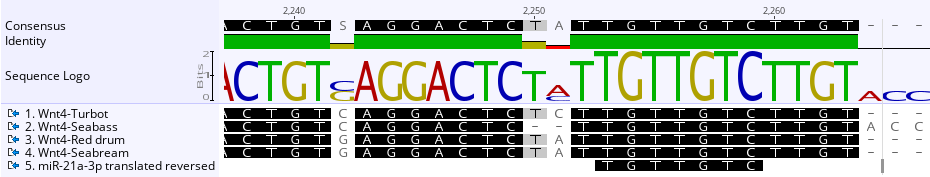

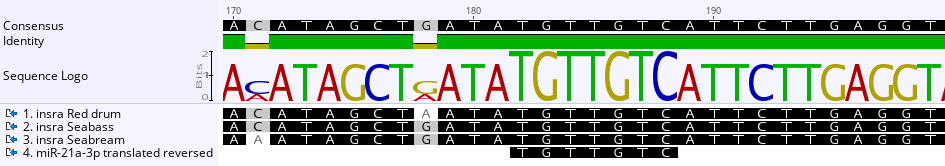

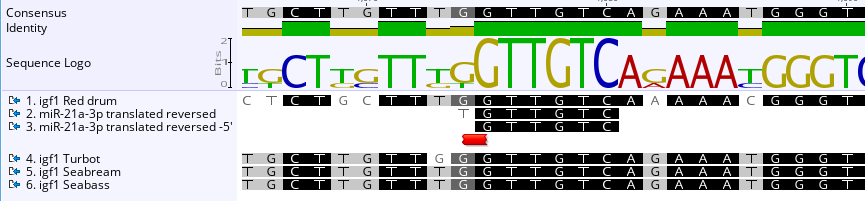

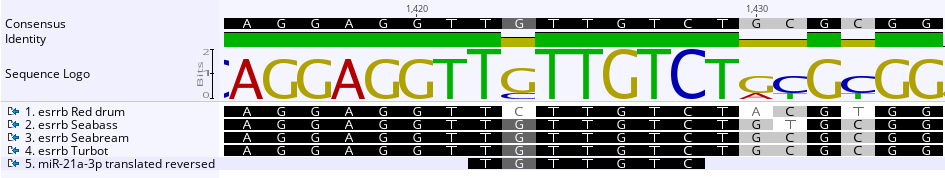

Alignment miR-125b-2/3-5p

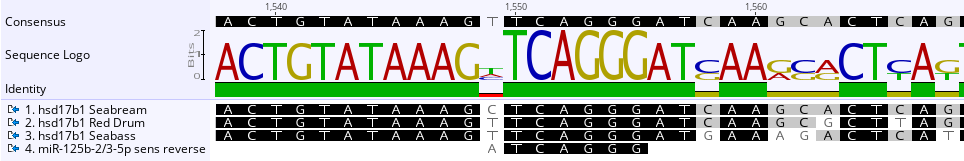

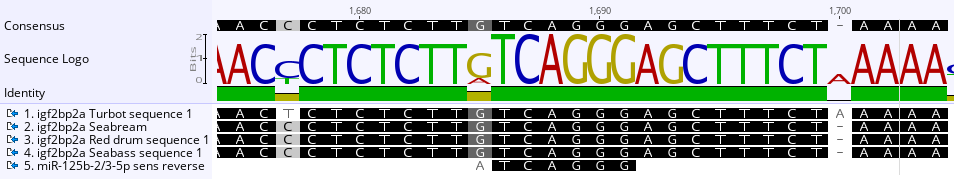

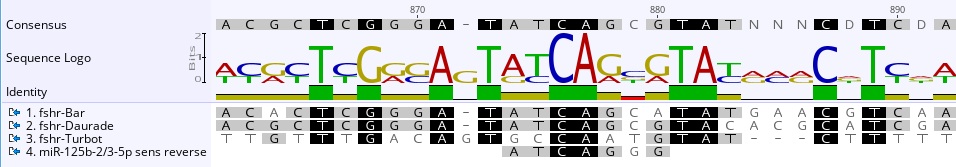
